# FK506-binding protein 13 expression is upregulated in interstitial lung disease and correlated with clinical severity: a potentially protective role

**DOI:** 10.1101/2019.12.15.858340

**Authors:** V Tat, EA Ayaub, A Ayoub, M Vierhout, S Naiel, MK Padwal, S Abed, O Mekhael, K Tandon, SD Revill, T Yousof, PS Bellaye, PS Kolb, A Dvorkin-Gheva, A Naqvi, JC Cutz, N Hambly, J Kato, M Vaughan, J Moss, MRJ Kolb, K Ask

## Abstract

Pulmonary fibrosis is a progressive lung disease characterized by myofibroblast accumulation and excessive extracellular matrix deposition. Endoplasmic reticulum (ER) stress initiates the unfolded protein response (UPR), a cellular stress response pathway that has been implicated in both inflammatory and fibrotic processes. Here, we sought to investigate the role of the 13 kDa FK506-binding protein (FKBP13), an ER stress-inducible molecular chaperone, in various forms of pulmonary fibrosis. We first characterized the gene and protein expression of FKBP13 in lung biopsy samples from 24 patients with idiopathic pulmonary fibrosis (IPF) and 17 control subjects. FKBP13 expression was found to be elevated in the fibrotic regions of IPF lung tissues, and within this cohort, was correlated with declining forced vital capacity and dyspnea severity. FKBP13 expression was also increased in lung biopsies of patients with hypersensitivity pneumonitis, rheumatoid arthritis, and sarcoidosis-associated interstitial lung disease. We next evaluated the role of this protein using FKBP13^-/-^ mice in a bleomycin model of pulmonary fibrosis. Animals were assessed for lung function and histopathology at different stages of lung injury including the inflammatory (Day 7), fibrotic (Day 21) and resolution (Day 50) phase. FKBP13^-/-^ mice showed increased infiltration of inflammatory cells and cytokines at Day 7, increased lung elastance and fibrosis at Day 21, and impaired resolution of fibrosis at Day 50. These changes were associated with an increased number of cells that stained positive for TUNEL and cleaved caspase 3 in the FKBP13^-/-^ lungs, indicating a heightened cellular sensitivity to bleomycin. Our findings suggest that FKBP13 is a potential biomarker for severity or progression of interstitial lung diseases, and that it has a biologically relevant role in protecting mice against bleomycin-induced injury, inflammation and fibrosis.

## INTRODUCTION

Pulmonary fibrosis results in the progressive scarring and stiffening of the lungs, eventually leading to respiratory failure [1]. Most cases are diagnosed as idiopathic pulmonary fibrosis (IPF), but secondary causes such as genetic susceptibility, environmental pollutants, medications and radiation exposure have been identified. Although a complete understanding of the cellular mechanisms involved in the pathogenesis of fibrotic lung disease remains elusive, accumulating evidence demonstrates that injury to the lung epithelium stimulates the accumulation of myofibroblasts which are responsible for the excessive deposition of extracellular matrix (ECM) components into the interstitium, causing decreased lung compliance and impaired gas exchange [1].

The endoplasmic reticulum (ER) is the site of protein biosynthesis and folding. Disruption of homeostasis in this organelle causes ER stress and activation of the unfolded protein response (UPR), which have been shown to contribute to the development of various fibrotic disorders including IPF [2, 3]. Activation of the UPR pathway in mammalian cells is thought to be triggered by the dissociation of Glucose-Regulated Protein 78 (GRP78) from three ER transmembrane proteins: inositol-requiring enzyme 1 (IRE1), activating transcription factor 6 (ATF6), and protein kinase RNA-like endoplasmic reticulum kinase (PERK). Activation of these pathways results in the upregulation of transcription factors and chaperones that increase the protein folding capacity of the ER and promote the degradation of misfolded or unfolded proteins [4]. Failure to restore ER homeostasis can lead to the initiation of apoptotic pathways through the transcription factor C/EBP-homologous protein (CHOP).

A distinct family of immunophilins termed the FK506-binding proteins (FKBPs) has been noted to play an important role in both connective tissue remodeling and the ER stress response [5, 6]. The mechanism by which FKBPs facilitate remodeling is thought to occur through their peptidyl– prolyl cis/trans–isomerase (PPIase) activity, which alters the peptidyl–prolyl bonds in proteins. FKBP65 is one such member of the family that is upregulated in lung tissue of IPF patients and that is necessary for collagen production by lung fibroblasts [5]. FKBP65 associates with GRP78 and the collagen-specific chaperone HSP47 during collagen assembly and its deficiency in mouse embryos results in altered collagen trafficking and aggregation of procollagen within the ER [7–9]. Here, we characterize the role of another one of the 15 members of FKBP family, the 13-kDa FK506-binding protein (FKBP13), in IPF patients and an experimental mouse model of bleomycin-induced lung fibrosis.

FKBP13 is localized to the lumen of the ER and is induced during ER stress [10–12]. It also shares 50% homology with other UPR chaperones GRP78 and GRP94, suggesting that FKBP13 itself also functions as a chaperone [11]. Boon et al. (2009) observed that FKBP13 gene expression was upregulated 5-fold in the lung tissue of patients with progressive IPF compared to those with stable disease [13]. Similar observations have been made in a cohort of patients with progressive hypersensitivity pneumonitis [14]. Unlike FKBP65, FKBP13 has lower activity towards post-translationally modified procollagen and does not appear to play a role in collagen folding [15]. Recently, it was shown that FKBP13 protects plasma cells from ER stress-induced apoptosis by promoting the degradation of misfolded immunoglobulins [16]. We postulated that the upregulation of FKBP13 in fibrotic lung tissue acts to protect cells against ER stress. However, as shown in our previous studies, its overall effect on the development of fibrosis depends on the specific cell types that are affected [17].

To further investigate the roles of FKBP13 in the pathogenesis of pulmonary fibrosis, we began by constructing a tissue microarray comprised of lung biopsies from IPF and control patients. This was used to investigate the expression pattern of FKBP13 at the protein and mRNA levels and to assess the association between FKBP13 expression and lung function in the patients. We then generated FKBP13-knockout (FKBP13^-/-^) mice and subjected them to a model of bleomycin-induced pulmonary fibrosis. In these experiments, we observed that FKBP13^-/-^ mice are more susceptible to injury by bleomycin, resulting in more severe pulmonary fibrosis at lower doses of the drug. These data suggest that FKBP13 plays a protective role in the pathogenesis of pulmonary fibrosis.

## Materials and Methods

### Human Lung Resected Tissue

All procedures using human tissues were approved by the Hamilton Integrated Research Ethics Board (11-3559 and 13-523-C). Formalin-fixed paraffin-embedded (FFPE) human lung tissue were obtained from the biobank for lung diseases at St. Joseph’s Hospital in Hamilton Ontario. IPF (n=30), hypersensitivity pneumonitis (n=5), rheumatoid arthritis (n=6) and sarcoidosis (n=6) cases were selected based on clinical history, radiographic appearance and a pattern of usual interstitial pneumonia or fibrotic granulomas as determined by trained molecular pathologists and radiologists. Non-involved tissue from lung cancer cases (n=17) was used as controls. Upon confirmation of a positive diagnosis, patient slides were scanned using the Olympus VS120 Slide Scanner and 0.6 mm diameter fibrotic and non-fibrotic cores were selected to be placed into a tissue microarray (TMA) block. Fibrotic regions were identified based on features of UIP including spatial heterogeneity in the parenchyma, architectural distortion and fibroblastic foci. In total, 316 cores were selected with at least 3 cores for each specified area. Forced vital capacity and Modified Medical Research Council (mMRC) dyspnea scores were obtained for the corresponding cases. Pulmonary function testing was conducted within a median of 48 days (range, 1-191 days) from the date of the biopsy.

### Histology

Tissue slides were stained with Hematoxylin and Eosin (H&E) for cellular analysis and tissue architectural analysis, Picrosirius Red (PSR) and Masson’s Trichrome for collagen analysis, anti-αSMA (Dako, Mississauga, ON, Canada, M0851) for the identification of myofibroblasts, GRP78 (Santa Cruz Biotechnology, N-20) and XBP1 to assess UPR activation, and anti-FKBP13 to visualize FKBP13 (R&D systems, Minneapolis, MN, MAB4356). Terminal deoxynucleotidyl transferase dUTP nick end labeling (TUNEL) was performed using the TACS® 2 *in situ* apoptosis detection kit (Trevigen, 4812-30-K). Stained slides were scanned with the Olympus VS120 Slide Scanner (Olympus, Waltham, MA) and analyzed with HALO Image Analysis Software (Indica Labs, Corrales, NM). A molecular pathologist (Dr. A. Naqvi) was consulted for histopathological analysis of FKBP13-expressing cell types.

### Gene expression analysis

RNA was extracted from FFPE tissues using the RNeasy FFPE kit (QIAGEN, Valencia, CA). RNA concentrations were measured using a NanoDrop® spectrophotometer and 3.3.0 software (NanoDrop Technologies, Wilmington, DE). The Nanostring nCounter platform (Nanostring Technologies, Seattle, WA, USA) was used to quantify the expression of FKBP13 in human FFPE samples as previously described [17]. Statistical analysis was performed using the nSolver Analysis Software v4.0 (Nanostring Technologies). Raw gene expression data was normalized to five housekeeping genes (*HPRT1, POLR2A, PPIA, TBP, TUBA1A*) that were selected by the built-in geNorm algorithm [18].

### Single-cell RNA-seq

Data preprocessed using the Cell Ranger pipeline (10x Genomics) were obtained from GSE122960. Out of 16 available samples, all 8 samples from donors and 4 samples from IPF patients were downloaded and used for further analysis. Post-processing was performed following the description provided by Reyfman et al. (2018) [19] using Seurat [20] package in R. t-distributed stochastic neighbor embedding (tSNE plot), expression plot and violin plots were created using Seurat package. Cell populations were defined by using the genes reported to be differentially expressed between the cell populations [19].

### Animal Experiments

All animal work was approved by the Animal Research Ethics Board of McMaster University (Hamilton, ON, Canada) under protocol number 12.02.06. Male C57BL6/J mice aged 10-12 weeks were bred and housed at the McMaster University Central Animal Facility (CAF, Hamilton, ON, Canada). Animals were kept on a 12-hour light/dark cycle at a controlled temperature of 20-25°C and ambient humidity of ∼50%. The animals were allowed access to food and water and were fed *ad libitum*.

### Generation of FKBP13-Deficient Mice by Gene Targeting

FKBP13-deficient animals were initially generated in 129SvEvBrd embryonic stem cells (Taconic Biosciences, NY), and chimeric mice were made on a C57BL/6J (Jackson laboratory) background to produce F2 homozygous mutants by Dr. Barbara Hendrickson (University of Chicago). Jackson Laboratory re-derived these mice in the C57BL/6J strain and they were further back-crossed for 7 generations onto a C57BL6/J background. Mice heterozygous (FKBP13^+/-^) and homozygous (FKBP13^-/-^) for the FKBP13 mutation were characterized by Southern blot analyses of tail genomic DNA. Northern blot analyses of RNA from WT (FKBP13^+/+^) brain, kidney, liver, and spleen revealed easily detectable 0.6 kB FKBP13 mRNA. However, no FKBP13 mRNA was detected in FKBP13^-/-^ mouse-derived tissues despite loading similar amounts of RNA for all samples, as judged by actin probe hybridization. Immunostaining for FKBP13 using an anti-FKBP13 antiserum (a gift from Dr. S. Burakoff, Dana-Faber Cancer Institute, Boston, MA) confirmed a lack of FKBP13 protein in the FKBP13^-/-^ lung tissue (**Figure 3A**).

**Figure 1.**
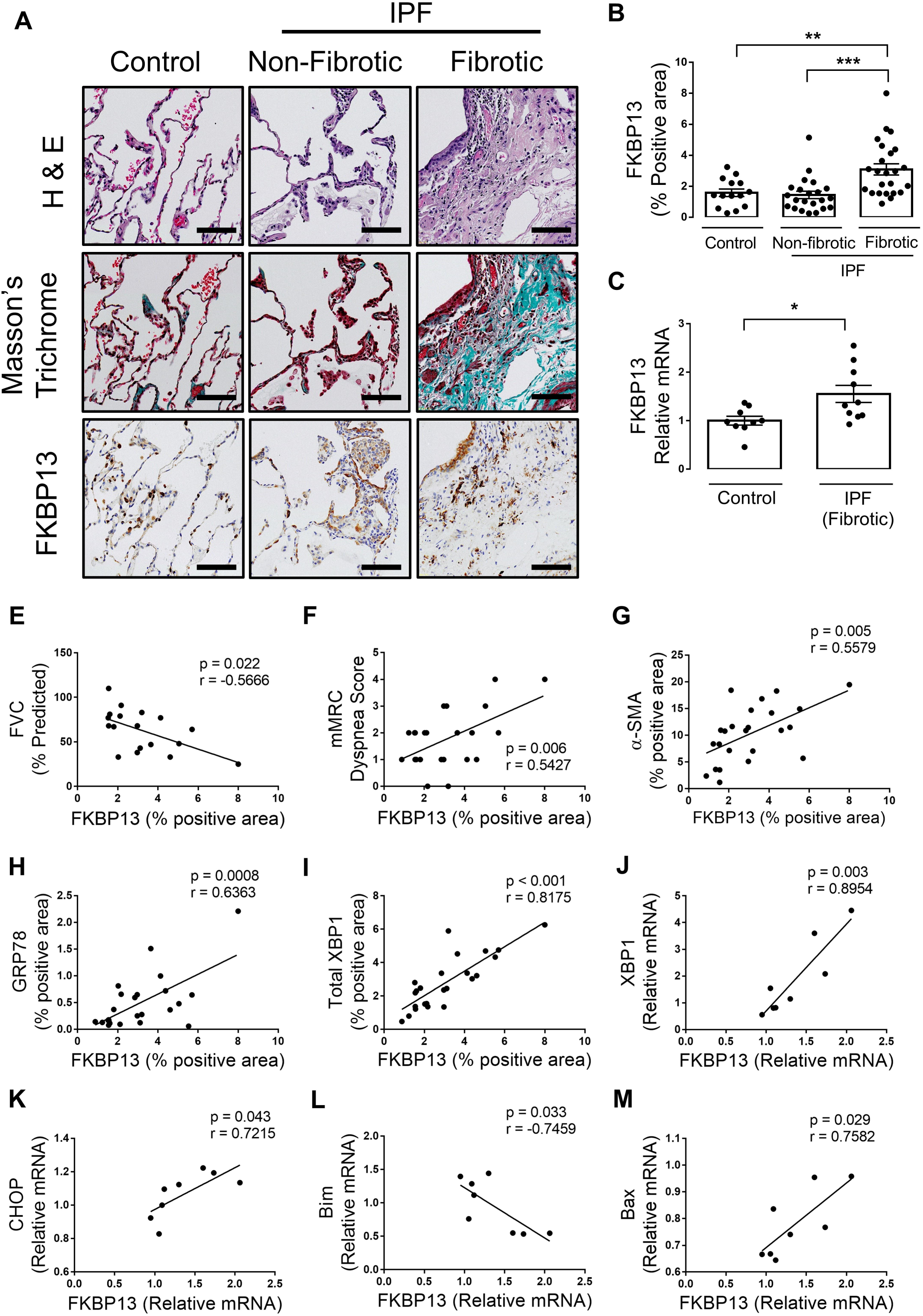
FKBP13 expression is elevated in fibrotic IPF tissue and is correlated with disease severity, ER stress and apoptosis markers. A tissue microarray containing fibrotic and non-fibrotic cores from 24 IPF patients and 17 controls was stained for FKBP13 and other markers by IHC and quantified by HALO image analysis software. NanoString nCounter® platform was used to assess the gene expression of UPR and apoptosis markers in the extracted cores. (A,B) Representative FKBP13 immunostaining images and quantification by HALO. Corresponding Masson’s Trichrome and H&E staining from serial sections are shown. Bar=100 µm. (C) Comparison of FKBP13 mRNA expression in fibrotic IPF and control tissue cores by Nanostring gene expression analysis. (D-H) Correlation of FKBP13-positive immunostained area with % predicted FVC, mMRC Dyspnea Score, α-SMA-, GRP78- and total XBP1-positive area. (I-L) Correlation of FKBP13 mRNA expression with XBP1, CHOP, Bim and Bax in the fibrotic regions of human IPF lung tissues. *, *p* < 0.05; **, p < 0.01 vs. Control.

**Figure 2.**
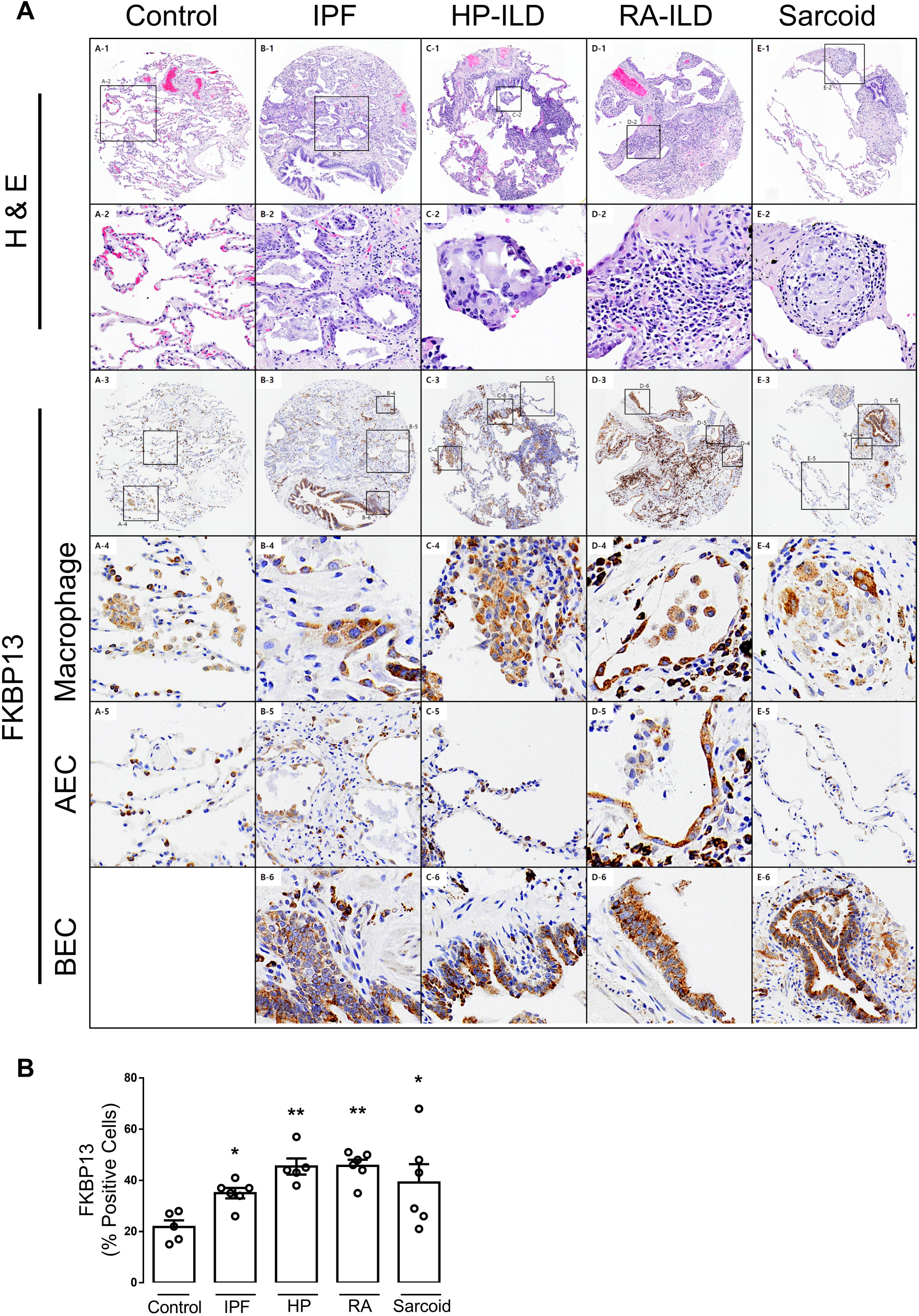
FKBP13 expression is elevated in other forms of interstitial lung disease. A second tissue microarray was constructed using lung biopsy cores from patients with interstitial lung disease secondary to hypersensitivity pneumonitis (HP), rheumatoid arthritis (RA) and sarcoidosis. A smaller, independent cohort of IPF patients was also included. (A) Representative H&E and FKBP13 immunostaining. Histologic hallmarks of each disease, as determined by a molecular pathologist, are shown (Row 2). Specific FKBP13-expressing cell types, including macrophages (Row 4), alveolar epithelial cells (AEC; Row 5) and bronchial epithelial cells (BEC; Row 6), were identified by the pathologist based on morphology. (B) Quantification of FKBP13 staining by HALO. *, *p* < 0.05; **, p < 0.01 vs. Control.

**Figure 3.**
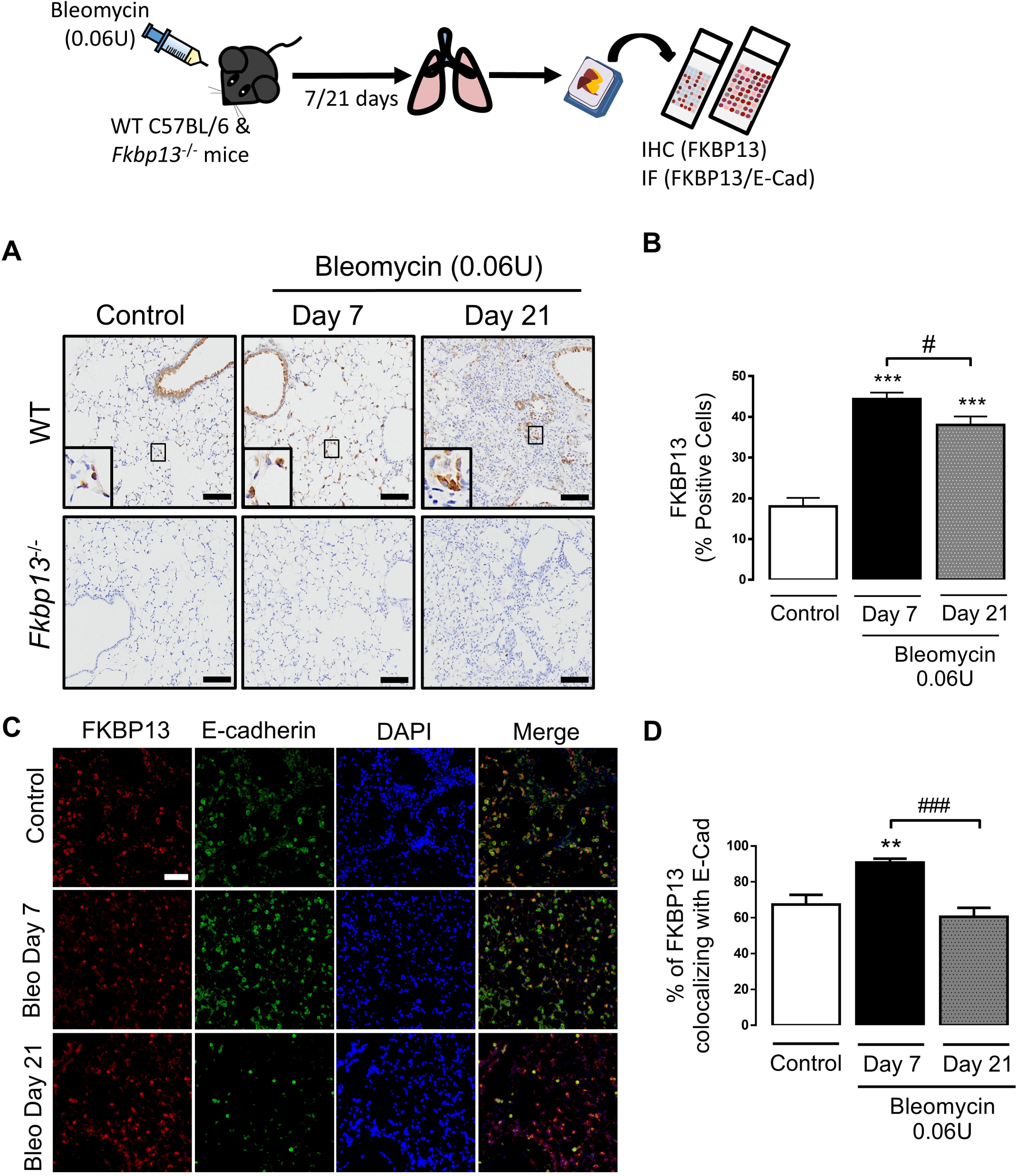
FKBP13 expression is induced by bleomycin in mice. Wildtype C57BL/6 mice and FKBP13^-/-^ (*n* = 4 per group) were treated with bleomycin (0.06U/mouse) and assessed after 7 and 21 days. Lung tissue was stained for FKPB13 by IHC and immunofluorescence. (A) Representative FKBP13 immunostaining images. Bar=100 µm. (B) Quantification of FKBP13 percent-stained area in WT mice by HALO image analysis software. No FKBP13 staining was detected in FKBP13^-/-^ lungs. *, *p* < 0.05 vs. untreated mice. (C) Dual immunofluorescence analysis confirmed the colocalization of FKBP13 (red) with the epithelial cell marker E-cadherin (green) in WT mice treated with bleomycin. DAPI was used as a nuclear marker. Bar=100 µm. (D) Percentage of FKBP13-positive area co-localizing with E-cadherin. **, *p* < 0.01; ***, *p <* 0.001 vs. untreated mice; #, *p* < 0.05; ###, *p* < 0.001 vs. Day 7.

### Bleomycin administration and collection of samples

Experimental pulmonary fibrosis was induced by intratracheal intubation of bleomycin (Hospira, NDC 61703-332-18) in isoflurane-anesthetized mice (MTC Pharmaceuticals, Canada) at 0.04 or 0.06U per mouse. Bleomycin was solubilized in 50µl sterile saline. Lung tissues and BALF were isolated and processed as described previously [17]. BALF differential cell counts were then performed by counting 300 leukocytes and using hemocytological procedures to classify the cells as neutrophils, macrophages, eosinophils or lymphocytes and multiplying the relative abundance of each cell type by the BALF total cell count. Mouse lung tissues were fixed and embedded in paraffin, and a TMA containing 135 cores of 1.5 mm diameter was generated using the TMA Master platform (3DHISTECH, Hungary). Three cores were obtained from each mouse lung, providing complete coverage of the lung area. Histological staining was performed on this TMA as described above. Semi-quantitative histological analysis of pulmonary fibrosis was performed through Ashcroft scoring as described previously [17].

### Immunofluorescence

Immunofluorescence staining of FKBP13 (R&D systems, Minneapolis, MN, MAB4356) and E-cadherin (Abcam, ab1416). Following deparaffinization and saturation of nonspecific sites with BSA (5%, 30 min), cells were incubated with primary antibodies overnight in a humidified chamber at 4°C. Conjugated secondary antibodies were used at a dilution of 1:2000. Slides were mounted in Prolong-gold with DAPI (ProLong® Gold antifade regent with DAPI, Life technologies, P36931). Images were taken using an epifluorescence microscope (Olympus IX81, Olympus, ON, Canada) and analyzed with MetaMorph® Image Analysis Software (Molecular Devices).

### Assessment of pulmonary mechanics

Lung function measurements, including pressure-volume loops and Quasi-static elastance were performed as described previously [17].

### Statistical analysis

Results were expressed as mean ± SEM. Two groups were compared with a two-tailed unpaired Student’s t-test. When more than two groups were compared, a one-way ANOVA followed by Newman-Keuls multiple comparison test was used. Statistical tests were employed using GraphPad Prism 7 (GraphPad Software, Inc). A p-value of <0.05 was considered statistically significant.

## RESULTS

### FKBP13 expression is increased in the fibrotic regions of IPF patients and is correlated with clinical severity

FKBP13 was previously shown to be upregulated at the mRNA level in patients with progressive IPF and other interstitial lung diseases [13]. Here, we aimed to characterize the expression of FKBP13 by immunohistochemistry (IHC) in the lungs of IPF patients. A tissue microarray was designed containing fibrotic and non-fibrotic cores taken from lung biopsies of 24 IPF patients and 17 control subjects. Demographic information and clinical characteristics of these patients are presented in **Table 1**. FKBP13 levels were higher in the fibrotic regions of IPF tissues compared to non-fibrotic regions and healthy control tissues (**Figure 1A,B**). Total RNA isolated from FFPE tissues was assessed for FKBP13 using Nanostring gene expression analysis. Consistent with the IHC findings, FKBP13 mRNA levels were elevated in the fibrotic tissues (**Figure 1C**). To determine the specific cellular subpopulations that express FKBP13, the open-access single-cell RNAseq dataset by Reyfman et al. (2018) was analyzed. Cells were clustered using tSNE and clusters were assigned to cell populations based on the genes reported as markers (**Figure S1A,B**). With the pooled data from lung biopsies of 4 IPF patients and 8 controls, FKBP13 was found to be ubiquitously expressed by all cellular subpopulations in the lung (**Figure S1C**) [19]. FKBP13 was noted to be significantly upregulated in the IPF plasma cell population.

**Table 1.** Baseline and functional characteristics of the IPF study population.

| Characteristic | IPF (N=24) |
| --- | --- |
| Age – years | 59.0±10.1 |
| Male sex – no. (%) | 17 (70.8) |
| Smoker (Former and Active) – no. (%) | 16 (66.6) |
| Disease Duration - years | 2.64±2.82 |
| FVC – % of predicted value | 62.9±23.2 |
| FEV<sub>1</sub> – % of predicted value | 64.5±20.3 |
| D<sub>LCO</sub> – % of predicted value | 42.8±10.9 |
| mMRC Dyspnea score* | 1.9±1.2 |
| Use of supplemental oxygen – no. (%) | 1 (4.2) |

\* Dyspnea was evaluated using the modified Medical Research Council (mMRC) scale. Scores range from 0 to 4, with higher scores indicating worse dyspnea.

Within the cohort of IPF patients, the expression of FKBP13 in the fibrotic cores was correlated with clinical parameters of disease severity. Specifically, we found a negative correlation with forced vital capacity (FVC) and a positive correlation with patient-reported dyspnea scores (**Figure 1D,E**). We further assessed the correlation of FKBP13 with other molecular markers of fibrosis, ER stress and apoptosis that have been linked to the development of IPF [21]. At the protein level, FKBP13 was positively correlated with the myofibroblast marker α-smooth muscle actin (αSMA) and the UPR markers GRP78 and total XBP1 (**Figure 1G-I**). This was corroborated at the mRNA level, with FKBP13 showing positive correlations with GRP78, total XBP1, CHOP and Bax, and a negative correlation with Bim (**Figure 1K-M**). In summary, higher levels of FKBP13 in fibrotic IPF tissues are associated with increased clinical severity and activation of UPR and apoptosis pathways, which may contribute to the disease pathogenesis.

We hypothesized that FKBP13 would also be upregulated in other forms of fibrotic lung disease. A second tissue microarray was constructed to assess the expression of FKBP13 in hypersensitivity pneumonitis, rheumatoid arthritis, and sarcoidosis-associated interstitial lung disease, in addition to an independent cohort of IPF patients. Baseline demographic and clinical data are shown in **Table 2**. Histopathological analysis of the FKBP13 staining by a molecular pathologist showed that FKBP13 appears primarily localized in the bronchial and alveolar epithelium, and in alveolar macrophages (**Figure 2A**). FKBP13 expression was found to be significantly elevated in the fibrotic regions of these tissues compared to control tissues (**Figure 2B**). These findings provide further support to potential role of FKBP13 in interstitial lung diseases.

**Table 2.** Baseline and functional characteristics of mixed interstitial lung disease study population.

| Characteristic | Idiopathic pulmonary fibrosis (N=6) | Hypersensitivity pneumonitis (N=5) | Rheumatoid arthritis (N=6) | Sarcoidosis (N=6) |
| --- | --- | --- | --- | --- |
| Age – years | 59.2±8.4 | 55.8±11.9 | 63.3±6.2 | 54.2±14.7 |
| Male sex – no. (%) | 2 (33.3) | 1 (20.0) | 5 (83.3) | 2 (33.3) |
| Smoker (Former and Active) – no. (%) | 2 (33.3) | 3 (60.0) | 5 (83.3) | 3 (50.0) |
| Disease Duration - years | 1.0±0.8 | 1.7±1.0 | 4.8±2.9 | 0.6±0.4 |
| FVC – % predicted | 57.2±17.0 | 66.8±17.8 | 61.5±6.1 | 65.3±17.2 |
| D<sub>LCO</sub> – % predicted | 64.0±17.3 | 43.0±13.0 | 32.2±5.9 | 52.5±11.0 |
| mMRC Dyspnea score* | 1.8±1.3 | 2.0±1.0 | 2.6±0.5 | 1.5±1.0 |

\* Dyspnea was evaluated using the modified Medical Research Council (mMRC) scale. Scores range from 0 to 4, with higher scores indicating worse dyspnea.

### FKBP13 expression is increased in the lung tissue of bleomycin-treated mice

Because of the upregulation of FKBP13 in various fibrotic lung diseases and its association with declining lung function and ER stress, we further explored the role of this protein in the development of pulmonary fibrosis by generating FKBP13 knockout (FKBP13^-/-^) mice. We induced pulmonary fibrosis using a single intratracheal 0.06U dose of bleomycin, and FKBP13 levels were assessed by IHC. In wildtype (WT) mice, FKBP13 was significantly upregulated at Day 7 and Day 21 post-bleomycin administration (**Figure 3A,B**). Based on histopathological analysis by a molecular pathologist, it was noted that FKBP13 was primarily localized in bronchial and alveolar epithelial cells, similar to the IPF tissues (**Figure 3A**). There was no FKBP13 staining present in FKBP13^-/-^ mice, providing validation for the knockout. Dual immunofluorescence analysis of WT lung tissues further confirmed the colocalization of FKBP13 with the epithelial cell marker E-cadherin (**Figure 3C**). Of note, the colocalization of FKBP13 and E-cadherin peaked at Day 7 (90.8±2.1%) before returning to baseline at Day 21 (**Figure 3D**).

### FKBP13 knockout increases susceptibility to bleomycin-induced pulmonary fibrosis

To determine if FKBP13^-/-^ mice developed a different fibrotic response to bleomycin administration compared to control mice, two doses of bleomycin, 0.04U or 0.06U/mouse, were delivered intratracheally to WT and FKBP13^-/-^ mice. Mice were then assessed for changes in lung function at Day 21 using a mechanical ventilator (Flexivent). At the lower dose of bleomycin (0.04U), WT mice were unaffected with respect to lung function and histopathology, while FKBP13^-/-^ mice displayed increased static lung elastance (**Figure 4A; Figure S2A**). At the higher dose of bleomycin (0.06U), both WT and FKPB13^-/-^ mice experienced a similar elevation in lung elastance at Day 21 (**Figure 4B; Figure S2B)**.

**Figure 4.**
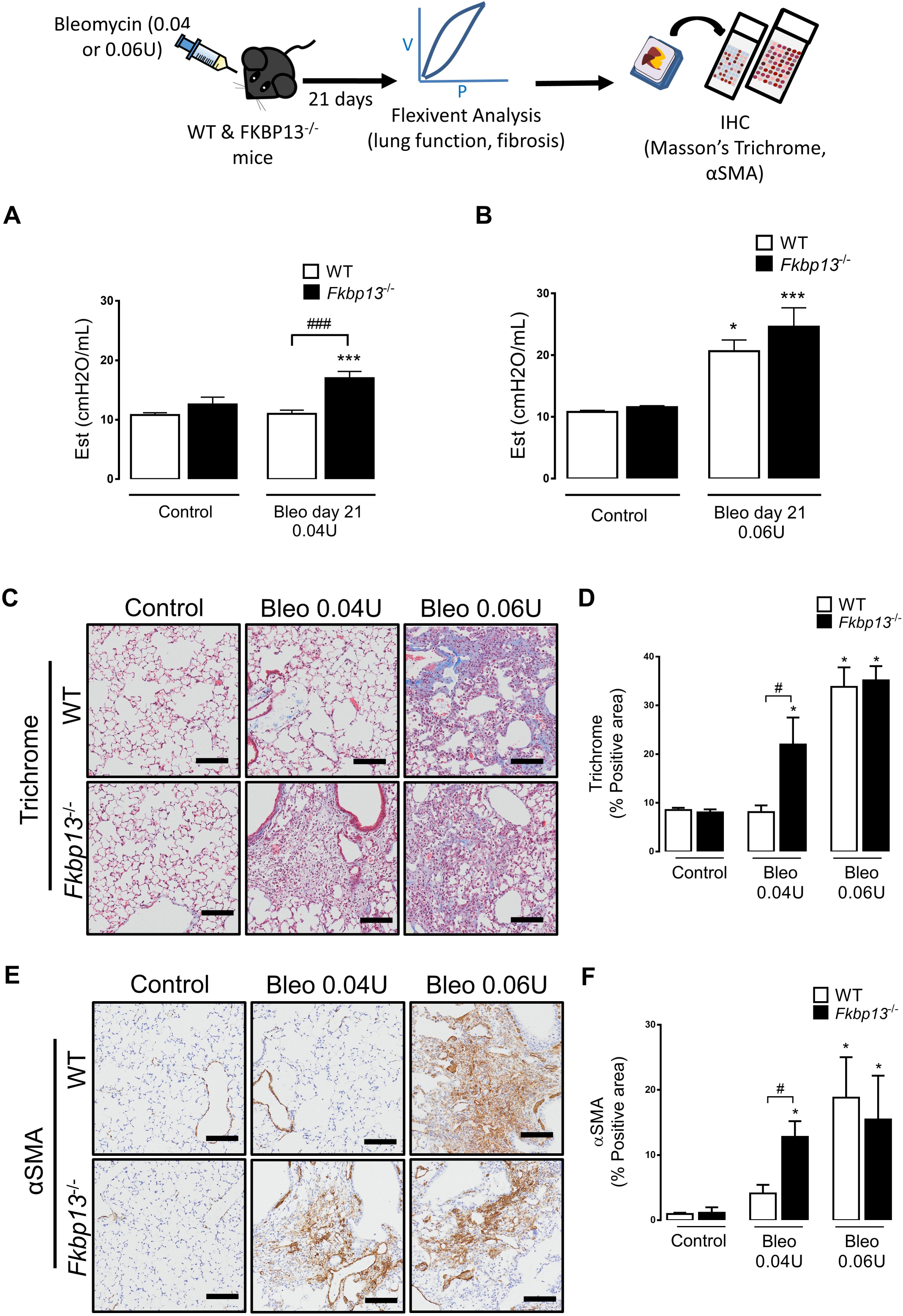
FKBP13^-/-^ mice are more susceptible to changes in lung function induced by a low dose of bleomycin. Wildtype and FKBP13^-/-^ mice (*n* = 6-8 per group) were treated with a low dose (0.04U/mouse) and high dose (0.06U/mouse) of bleomycin and lung function, as well as fibrosis, was assessed after 21 days. (A,B) Elastance measurements were derived from the PV loops (**Figure S2**). FKBP13^-/-^ mice are more susceptible to fibrosis induced by a low dose of bleomycin. (C,D) Masson’s Trichrome staining to assess collagen deposition and quantification of Trichrome-positive area by HALO image analysis software. Bar=100 µm. (E,F) Alpha smooth muscle actin immunostaining to assess myofibroblast accumulation and quantification by HALO. Bar=100 µm. *, *p* < 0.05; ***, *p* < 0.001 vs. untreated mice. #, *p* < 0.05; ###, *p* < 0.001 between genotypes.

To determine whether the higher elastance measurement of the FKBP13^-/-^ lungs at Day 21 correlated with an increase in fibrogenesis, histopathological analysis of the lung tissues was performed. Consistent with the lung stiffness measurements, the lower dose of bleomycin led to increased collagen deposition in the lung parenchyma of the FKBP13^-/-^ mice, while WT mice were unaffected (**Figure 4C,D**). The fibrotic lung parenchyma of FKBP13^-/-^ mice also showed increased αSMA immunostaining compared to WT mice, indicating an accumulation of myofibroblasts (**Figures 4E,F**). At the higher bleomycin dose, the extent of fibrosis and myofibroblast accumulation was relatively similar in both strains at Day 21. Taken together, these results show that FKBP13^-/-^ mice are more susceptible to bleomycin-induced lung injury, suggesting that FKBP13 is involved in processes that protect against either injury or fibrogenesis.

### FKBP13 deficiency increases lung inflammation after bleomycin administration

To determine whether the exaggerated fibrotic processes in the FKBP13^-/-^ mice are due in part to their susceptibility towards bleomycin-induced pulmonary inflammation, we assessed the inflammatory profile of the BALF collected from the mice, which included total cell infiltrates, cell differentials, TGF-β1 and IL-6 levels. This was done both at the peak of the inflammatory phase (Day 7) and the fibrotic phase (Day 21) after bleomycin instillation. At Day 7, the lower dose of bleomycin (0.04U) led to an influx of inflammatory cells in both WT and FKBP13^-/-^ mice compared to their respective controls; however, the FKBP13^-/-^ mice showed a two-fold greater influx over the WT mice (**Figure 5A**). FKBP13^-/-^ mice had increased infiltration of macrophages, neutrophils and lymphocytes at Day 7 compared to WT mice (**Figure 5B-D**). There were no differences in the amount and phenotype of inflammatory cells at Day 21 post-bleomycin administration. Levels of the pro-inflammatory cytokine IL-6 were elevated in both strains at Day 7, but to a 2-fold greater extent in FKBP13^-/-^ compared to WT lungs (**Figure 5E**). Levels of the pro-fibrotic cytokine TGF-β1 were also higher in FKBP13^-/-^ mice compared to WT mice at Day 7 (**Figure 5F**). At Day 21, the level of TGF-β1 was below detection. This data demonstrates that although both strains experience a pro-inflammatory response at Day 7, the response is intensified in the FKBP13^-/-^ mice.

**Figure 5.**
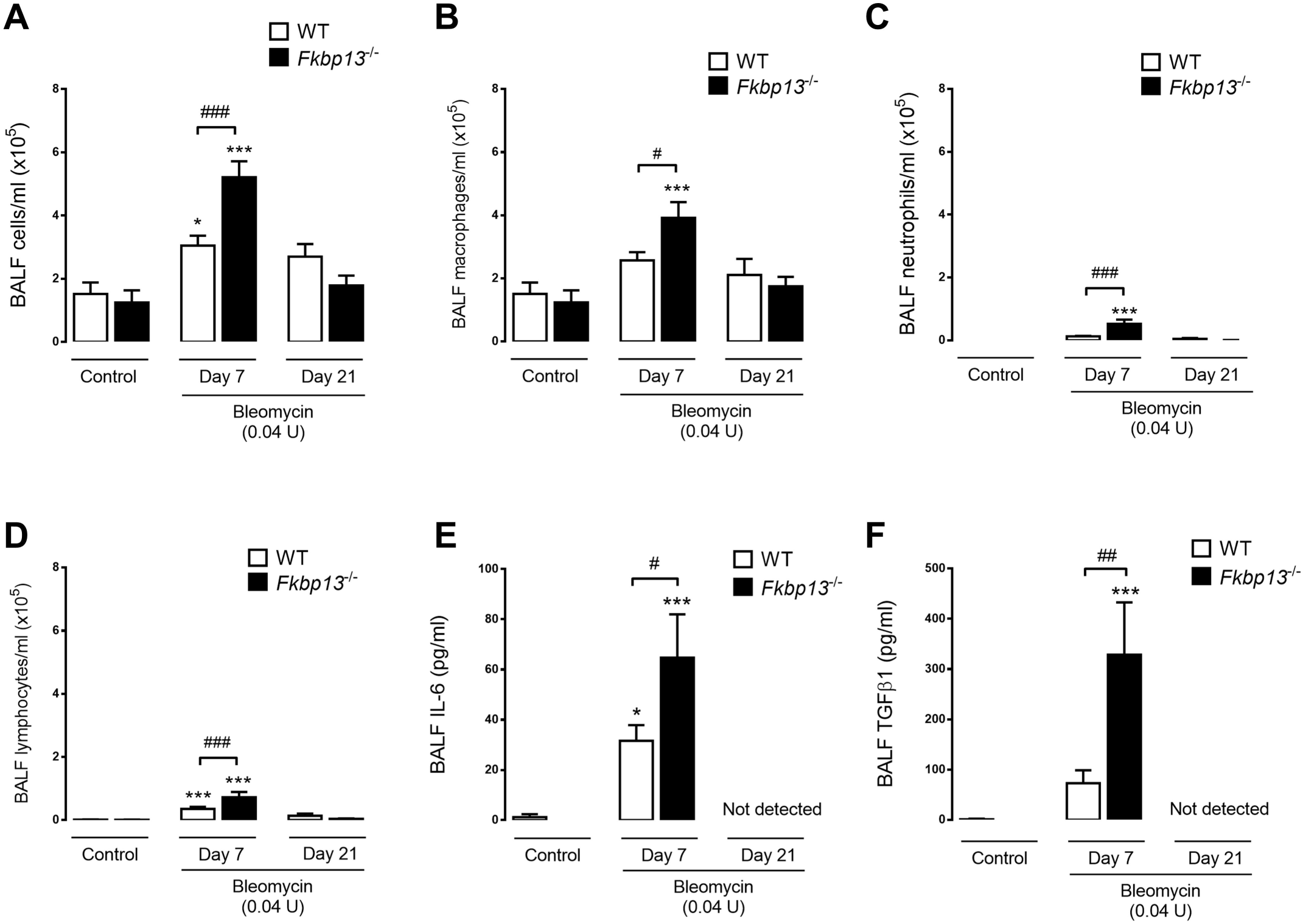
FKBP13^-/-^ mice have enhanced cellular infiltration and profibrotic cytokine signaling in response to a low dose of bleomycin. Wildtype and FKBP13^-/-^ mice (*n* = 6-8 per group) were treated with the low dose (0.04U/mouse) of bleomycin and BALF was collected at Day 7 to assess their inflammatory profiles. BALF cells were stained with Wright-Giemsa and the total cells (A), macrophages (B), neutrophils (C), and lymphocytes (D) were counted. (E,F) IL-6 and TGF-β1 levels in the BALF were assessed by ELISA.

### FKBP13^-/-^ mice are more susceptible to bleomycin-induced apoptosis

Epithelial injury and apoptosis are recognized as precipitating events in the development of pulmonary fibrosis [23]. To assess apoptosis in the bleomycin-treated mice, lung tissues were stained for TUNEL and cleaved caspase 3. At Day 7 following administration of the low dose of bleomycin, FKBP13^-/-^ mice demonstrated a significant increase in the number of TUNEL-positive cells (**Figure 6A,B**). Higher numbers of apoptotic cells were also seen in the knockout lung tissues at Day 21 by cleaved caspase 3 immunostaining (**Figure 6C,D**). Histopathological analysis suggested that type II alveolar epithelial cells were the predominant cell type affected.

**Figure 6.**
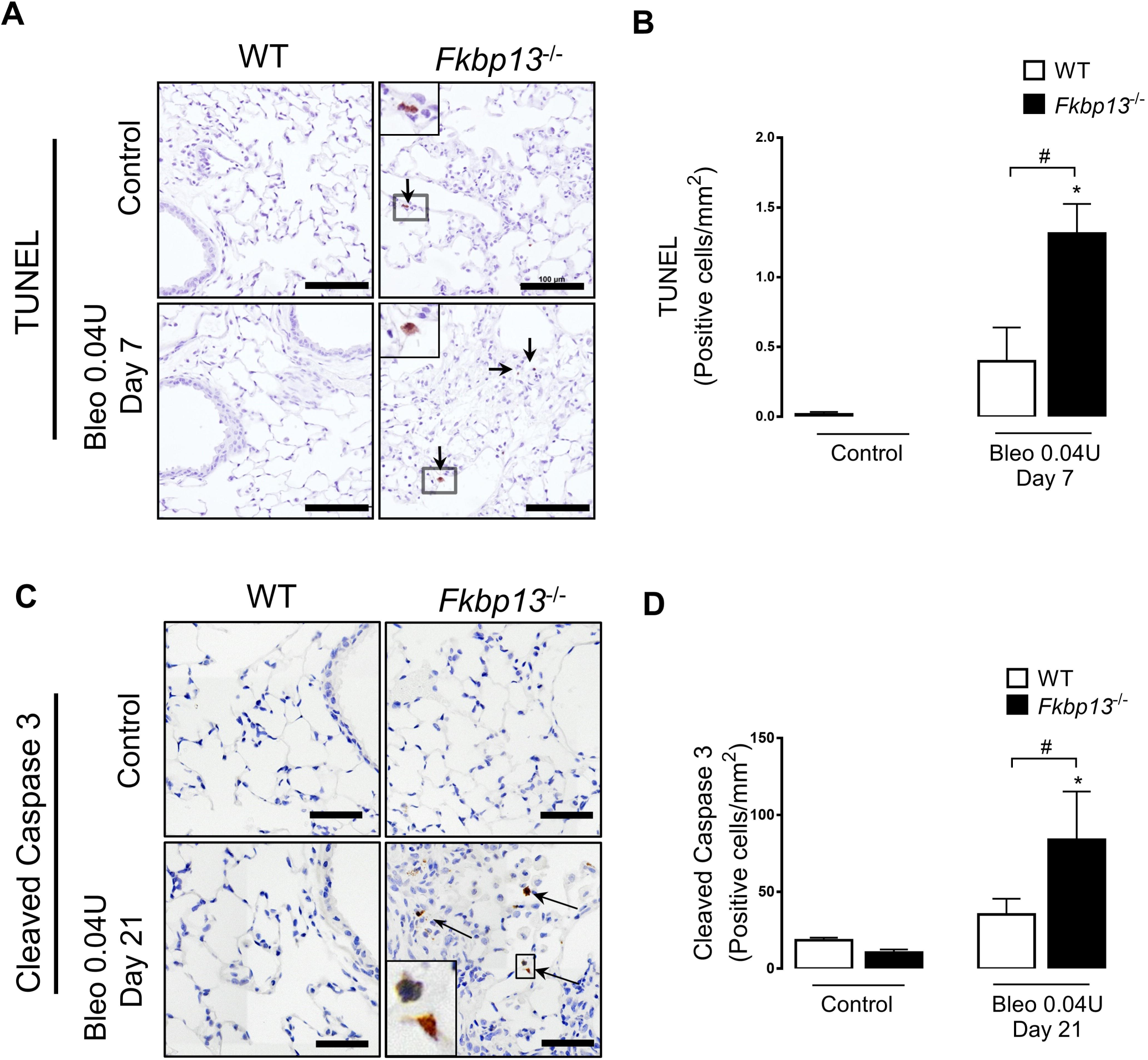
FKBP13^-/-^ lungs have increased levels of UPR activation and apoptotic cells. Wildtype and FKBP13^-/-^ mice (*n* = 6-8 per group) were treated with the low dose (0.04U/mouse) of bleomycin and lung tissues were assessed for UPR and apoptosis markers using IHC, in situ hybridization and Western blot. (A,B) TUNEL staining and quantification of positive cells at Day 7 by HALO image analysis software. Bar=100 µm. (C,D) Cleaved caspase 3 immunostaining and quantification of positive cells at Day 21 by HALO image analysis software. Bar=100 µm. *, *p* < 0.05 vs. untreated mice. #, *p* < 0.05 between genotypes.

### Resolution of bleomycin-induced pulmonary fibrosis is impaired in FKBP13^-/-^ mice

A characteristic of the bleomycin model is the reversibility of the fibrosis in the afflicted animals [22]. To determine if FKBP13 deficiency affects this process, lung stiffness and fibrosis measurements were made at 50 days following the administration of high dose (0.06U) bleomycin. As reported earlier, this dose causes both strains develop a similar severity of fibrosis at Day 21 (**Figure 4**). At Day 50, lung stiffness in the WT mice returned to baseline, while remaining elevated in the FKBP13^-/-^ mice (**Figure 7A,B**). Fibrotic tissue area also persisted in the FKBP13^-/-^ mice while normalizing in the WT mice (**Figure 7C,D**), suggesting that FKBP13 deficiency impairs fibrosis resolution.

**Figure 7.**
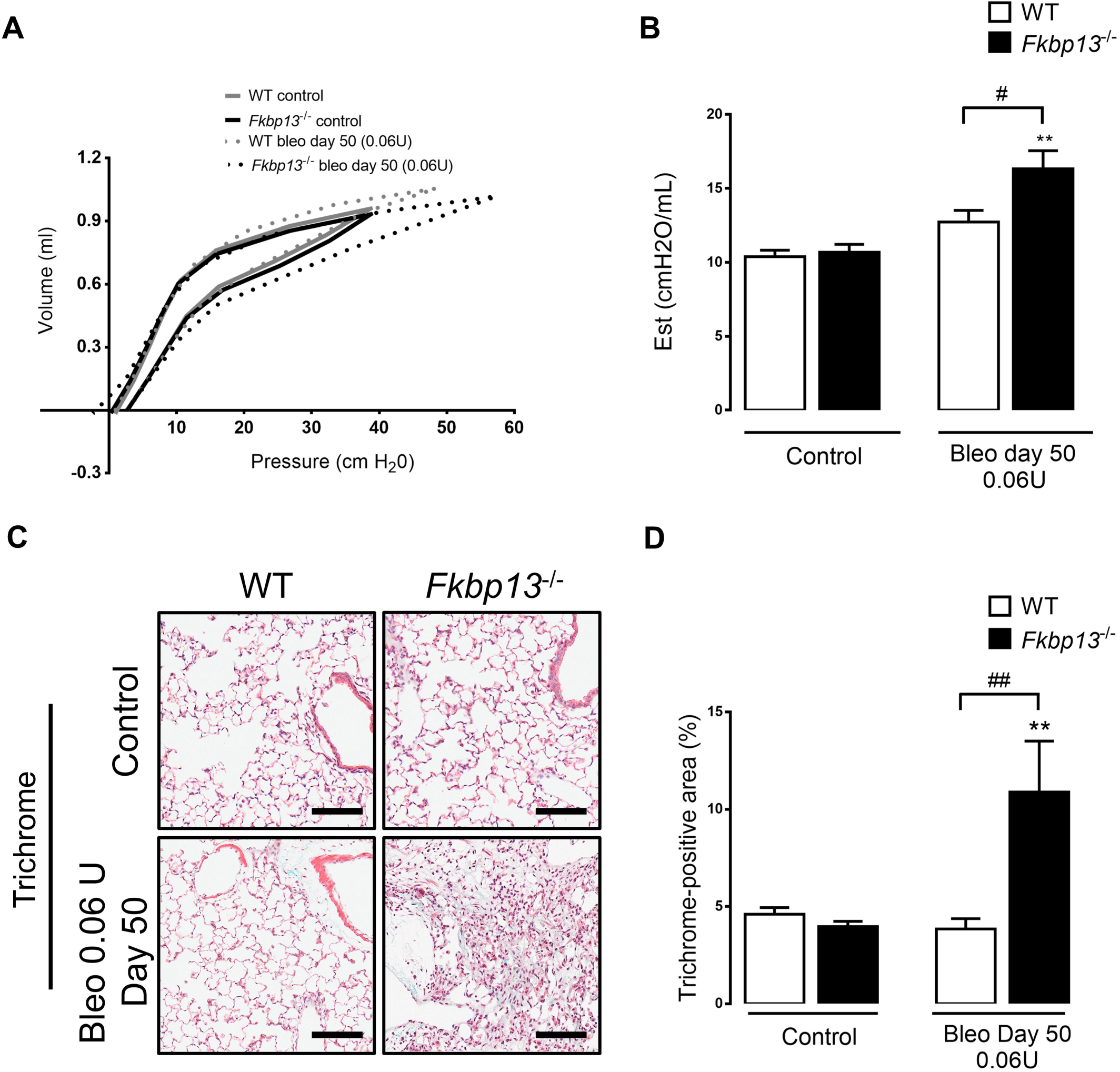
FKBP13^-/-^ mice have impaired resolution of bleomycin-induced pulmonary fibrosis. WT and FKBP13^-/-^ mice (*n* = 4-6 per group) were treated with high dose bleomycin and assessed for changes in lung function and fibrosis at Day 50. (A,B) Pressure-volume loops as assessed by flexiVent® and derivation of elastance. (C,D) Masson’s Trichrome staining of lung tissues for assessment of fibrosis and quantification of Trichrome-positive area by HALO. Bar=100 µm. **, *p* < 0.01 vs. untreated mice. #, *p* < 0.05; ##, *p* < 0.01 between genotypes.

## DISCUSSION

In this study, we investigated the role of the ER resident molecular chaperone, FKBP13, in patients diagnosed with IPF and in an *in vivo* system which models the pathogenesis of pulmonary fibrosis. Previously, FKBP13 has been identified as a gene that is upregulated in the lung tissue of patients with progressive IPF, as defined by a ≥10% decline in FVC and ≥15% decline in DL_CO_ over the 12 months following the biopsy [13]. Furthermore, FKBP13 expression was found to be elevated in a cohort of patients with rapidly progressive chronic hypersensitivity pneumonitis, a disease that is also characterized by inflammation and progressive fibrosis [14]. In our present study, we corroborate these findings at the protein level, with higher expression of FKBP13 detected in the fibrotic lung tissue of IPF, hypersensitivity pneumonitis, rheumatoid arthritis and sarcoidosis patients compared to non-fibrotic regions and control tissues. We also report that FKBP13 expression in IPF fibrotic lung tissue correlates with parameters of disease severity such as FVC and patient-reported dyspnea scores, suggesting that it could be useful as a biomarker for disease severity and progression. Our studies in transgenic mice confirmed a strong pathophysiological relevance of this molecule in all stages of lung injury including the inflammatory, fibrotic and resolution phases. We demonstrate that complete knockout of FKBP13 in mice results in increased susceptibility to bleomycin-induced pulmonary fibrosis and impaired resolution of fibrosis.

Bleomycin is known to induce overall ER stress levels in the mouse lung [17, 24, 25], but due to the heterogeneous nature of this organ, an important question that remains is determining the specific cellular subpopulations that experience this ER stress and their resulting contribution to fibrotic remodeling. Activation of the UPR in response to ER stress initially leads to an adaptive adjustment of protein folding capacity through the upregulation of ER resident molecular chaperones, attenuation of protein synthesis, degradation of misfolded or unfolded proteins, and physical expansion of the ER [26]. Failure of these cellular programs to restore ER homeostasis results in the induction of apoptosis through transcription factors such as CHOP. As a component of the adaptive phase of the UPR, molecular chaperones such as FKBP13 are important for promoting cell survival through the *de novo* folding and assembly of proteins, refolding of misfolded proteins, and facilitation of proteolytic degradation^16^.

Each cellular subpopulation of the lung has a potential role in either promoting or protecting against the fibrotic remodeling process. Because of the pro-survival role of chaperones, the contribution of a molecular chaperone to the pathogenesis of IPF therefore depends on its net effect on all cell types in which it is expressed. Profibrotic chaperones include those that maintain the function of matrix- and cytokine-producing cell types such as fibroblasts and macrophages. For example, calreticulin, FKBP65, GRP78 and HSP47 are ER resident chaperones that maintain myofibroblast function by regulating Type I collagen biosynthesis and incorporation into the ECM of fibrotic tissues [5, 27, 28]. We have previously shown that GRP78 also contributes to the persistence of M2-like macrophages in fibrotic lung tissue by protecting these cells from undergoing CHOP-mediated apoptosis [17]. By contrast, antifibrotic chaperones include those that promote the maintenance of a normal epithelium in response to pulmonary insults, and that restrict the function of profibrotic cell types. For example, HSP70 has been shown to prevent epithelial cells from undergoing TGF-β1-induced epithelial-mesenchymal transition, and its overexpression in mice protects against bleomycin-induced pulmonary fibrosis [29]. We found, based on single-cell RNAseq analysis, that FKBP13 mRNA is present in all pulmonary cell types, and thus its loss can have varying degrees of impact on fibrogenesis depending on the cell type affected. Histopathological analysis of FKBP13 staining in fibrotic lung tissue revealed predominant expression of this protein in epithelial cells and alveolar macrophages. Here, we report evidence that the complete loss of FKBP13 has a net fibrotic effect on the development of pulmonary fibrosis. We further explored the impact of this loss on epithelial cell injury and cellular infiltrates into the lung tissue, both of which have been shown to correlate with the extent and severity of fibrosis in the bleomycin model [30, 31].

Disturbance of the bronchial and alveolar epithelium through apoptosis has been observed in human IPF lung tissue, and is considered to be an initial event in acute lung injury that leads to the subsequent fibrotic response [21, 23, 32]. There is now increasing evidence that prolonged ER stress can induce these apoptotic processes in alveolar epithelial cells [33–35]. In human IPF lung tissues, the levels of FKBP13 in fibrotic regions correlated with various markers of ER stress and apoptosis, indicating a potential involvement of this chaperone in these pathways. We observed here that a deficiency in FKBP13 increased the sensitivity of mice to low doses of bleomycin-induced pulmonary fibrosis, suggesting that the epithelium is more susceptible to injury. Consistent with this, we observed an increase in apoptotic epithelial cells in the FKBP13^-/-^ lung tissue at both Day 7 and Day 21 post-bleomycin administration. Given its role as a molecular chaperone, we postulate that FKBP13 serves to protect epithelial cells from ER stress-mediated apoptosis by maintaining ER homeostasis. This is consistent with the findings of Jeong et al. (2017), who found that FKBP13 is induced by ER stress in plasma cells and acts to limit the production of immunoglobulins in the ER, thereby preventing the cells from undergoing CHOP-mediated apoptosis [16]. In our model, the loss of FKBP13 may impair the ability of epithelial cells to adapt to ER stress, priming them for more apoptosis in response to an exogenous insult such as bleomycin. This is consistent with previous studies in which a chaperone deficiency resulted in constitutive activation of the UPR because of higher basal levels of ER stress [17, 36]. Although our knockout experiments suggest that FKBP13 is antifibrotic, its upregulation in both human and mouse fibrotic lung tissue could be a physiological response to cell injury and ER stress. A defect in the chaperone function of FKBP13 or prolonged occupation of its binding site could presumably lead to a similar phenotype as the FKBP13^-/-^ mice.

The immune cells that infiltrate the lung tissue during bleomycin injury lead to the generation of pro-inflammatory cytokines, which is followed by a switch at around Day 9 to pro-fibrotic signaling. To investigate whether the exacerbated fibrotic response was in part due to an increased cellular infiltration of FKBP13^-/-^ lungs, we examined the inflammatory profile of the BALF. In response to the low dose of bleomycin, both strains displayed cellular infiltration and significantly elevated IL-6 at Day 7 over their respective controls, although the levels in the FKBP13^-/-^ mice were two-fold greater than the WT mice. We have previously shown that IL-6 potentiates the pro-fibrotic phenotype of M2 macrophages, driving the fibrosis resulting from bleomycin injury [37]. TGF-β1 levels were also elevated in the knockout mice at Day 7, and is known to stimulate myofibroblast proliferation and activation, as indicated by the increased α-SMA staining. In addition to the more robust cellular infiltration seen in the knockout, the loss of FKBP13 may alter the intrinsic functioning of the responding immune cells. Our single-cell RNAseq analysis found elevated FKBP13 expression in plasma cells while histopathological analysis revealed positive staining of the protein in alveolar macrophages. The elevated cytokine levels in FKBP13^-/-^ mice may be the result of unopposed cytokine production by these cells. It has been shown in plasma cells that FKBP13 acts to restrict IgA production in order to maintain ER homeostasis and promote cell survival [16], and therefore a similar mechanism may be involved in regulating the output of cytokines. Furthermore, the milder but significant acute inflammation seen in the WT mice suggests that these animals still experienced some degree of lung injury in response to bleomycin but were able to resolve the injury without progressing to the fibrotic stage. The increased resistance of epithelial cells to apoptosis and the subdued immune response may both contribute to the protective effect of FKBP13 in WT mice.

The bleomycin model is characterized by an ability of the afflicted mice to recover and undergo resolution of fibrosis. Several mechanisms of pulmonary fibrosis resolution have been proposed in the literature, including the degradation of the fibrotic ECM, removal of myofibroblasts, re-epithelialization, and removal of inflammatory cells such as profibrotic macrophages [22]. At the higher bleomycin dose of 0.06U, both WT and FKBP13^-/-^ mice displayed a similar increase in lung elastance measurements at Day 21, implying that although the FKBP13^-/-^ mice are more sensitive to bleomycin-induced pulmonary fibrosis, the extent of fibrosis approaches the same plateau as the bleomycin dose increases. At Day 50, lung elastance returned to baseline in the WT but remained elevated in the FKBP13^-/-^ mice, suggesting that the loss of FKBP13 impairs the resolution of fibrosis in the bleomycin model.

FK506 (tacrolimus) and rapamycin (sirolimus) are immunosuppressive drugs that bind to members of the FKBP family, including FKBP13. Several clinical and animal studies have yielded conflicting results regarding the effect of these drugs on pulmonary fibrosis. Inhibition of FKBP13 chaperone activity by these drugs could result in a similar phenotype as FKBP13 deficiency, with a predisposition to epithelial injury and intensified inflammation [16]. Our findings provide a potential explanation for the interstitial pneumonitis and focal pulmonary fibrosis that has been reported in transplant patients being treated with these immunosuppressants [38, 39]. FK506 has shown both beneficial and adverse effects on connective tissue disease-associated interstitial lung disease, with improvements seen in myositis patients [40–42] and pulmonary injury reported in patients with rheumatoid arthritis [43, 44]. The effect of rapamycin on bleomycin-induced pulmonary fibrosis is also inconclusive, with some studies suggesting that it inhibits the development of fibrosis [45] and others suggesting that it worsens the disease [46]. These studies largely attributed the effects of rapamycin to its interaction with FKBP12 and the subsequent inhibition of the PI3K/AKT/mTOR pathway [45, 47], but inhibition of the other FKBPs such as FKBP13 is likely involved in its effects. The overall effect of medications such as rapamycin and FK506 on pulmonary fibrosis likely depends its distribution across the various cellular compartments, its relative affinity to the FKBPs, and the etiology of the disease.

A limitation of our study is the lack of prognostic data on the IPF patients that were enrolled. Pulmonary function tests were conducted within a median of 48 days from the date of biopsy. Additional data on patient survival and serial pulmonary function testing results may aid in further study of the prognostic relevance of FKBP13. To further investigate the pathophysiological role of FKBP13 in pulmonary fibrosis, next steps would involve the generation of tissue-specific knockouts in the epithelial or myeloid lineage to stratify the effects on epithelial injury and inflammation.

In conclusion, our data suggests that FKBP13 is upregulated and plays a protective role in the pathogenesis of pulmonary fibrosis. Mice lacking this chaperone are more susceptible to injury from lower doses of bleomycin. Because FKBP13 is predominantly expressed in lung epithelial cells, we postulate that the loss of FKBP13 reduces the ability these cells to respond to ER stress, priming them for more injury and apoptosis in response to pulmonary insults. In human IPF lung tissues, levels of FKBP13 are associated with clinical parameters of disease severity. While unlikely to be a pharmacological target because of its protective role, next steps should involve validation of FKBP13 as a potential biomarker for the progression of fibrotic interstitial lung diseases.

## ACKNOWLEDGEMENTS

We sincerely thank the central animal facility and McMaster Immunology Research Centre’s Core Histology Facility (Mary Jo Smith) for their exceptional technical training and service; Jane Ann Smith (McMaster University, ON, Canada) for her exceptional technical assistance with the experimental animal work; Jennifer Wattie (St. Joseph’s Healthcare Hamilton, ON, Canada) for technical help in performing flexiVent lung function assessments; Christine Mader (Farncombe Metagenomics Facility, McMaster University, ON, Canada) for help with NanoString experiments; and Pavithra Parthasarathy, James Murphy, Dr. Carl Richards, Dr. Martin Stämpfli, Dr. Jeffrey Dickhout and Dr Jack Gauldie for their support and constructive discussions. This study was supported by funding from the Canadian Pulmonary Fibrosis Foundation, Ontario Thoracic Society, CFI awarded to KA. KA reports sponsored collaborative research projects with Genoa, Windward, Actelion, Gilead, Patara, Boehringer Ingelheim, Synairgen, Alkermes, GSK, Pharmaxis, Indalo, Unity and Avalyn, outside of the submitted work. EAA was funded by the Canadian Institute of Health Research (CIHR; Grant No. 140358). MK reports grants from CIHR; grants and personal fees from Roche, Boehringer Ingelheim and Prometic, grants from Actelion, Respivert and Alkermes, and personal fees from Genoa, Indalo and Third Pole, outside the submitted work. JM, MV and JK were supported by the NIH Intramural Research Program.

## Notes

**Conflict of interest:** No conflict of interests were declared

## REFERENCES

[1] Lederer DJ, Martinez FJ. Idiopathic Pulmonary Fibrosis. N. Engl. J. Med. 2018;378:1811–23.

[2] Wei J, Rahman S, Ayaub EA, Dickhout JG, Ask K. Protein Misfolding and Endoplasmic Reticulum Stress in Chronic Lung Disease. Chest. 2013;143:1098–105.

[3] Kropski JA, Blackwell TS. Endoplasmic reticulum stress in the pathogenesis of fibrotic disease. J. Clin. Invest. 2018;128:64–73.

[4] Yoshida H, Matsui T, Yamamoto A, Okada T, Mori K. XBP1 mRNA Is Induced by ATF6 and Spliced by IRE1 in Response to ER Stress to Produce a Highly Active Transcription Factor. Cell. 2001;107:881–91.

[5] Staab-Weijnitz CA, Fernandez IE, Knüppel L, Maul J, Heinzelmann K, Juan-Guardela BM, et al. FK506-Binding Protein 10, a Potential Novel Drug Target for Idiopathic Pulmonary Fibrosis. Am. J. Respir. Crit. Care Med. 2015;192:455–67.

[6] Liu Z, Cai H, Zhu H, Toque H, Zhao N, Qiu C, et al. Protein kinase RNA-like endoplasmic reticulum kinase (PERK)/calcineurin signaling is a novel pathway regulating intracellular calcium accumulation which might be involved in ventricular arrhythmias in diabetic cardiomyopathy. Cell. Signal. 2014;26:2591–600.

[7] Duran I, Martin JH, Weis MA, Krejci P, Konik P, Li B, et al. A Chaperone Complex Formed by HSP47, FKBP65 and BiP Modulates Telopeptide Lysyl Hydroxylation of Type I Procollagen. J. Bone Miner. Res. Off. J. Am. Soc. Bone Miner. Res. 2017;32:1309–19.

[8] Ishikawa Y, Holden P, Bächinger HP. Heat shock protein 47 and 65-kDa FK506-binding protein weakly but synergistically interact during collagen folding in the endoplasmic reticulum. J. Biol. Chem. 2017;292:17216–24.

[9] Lietman CD, Rajagopal A, Homan EP, Munivez E, Jiang M-M, Bertin TK, et al. Connective tissue alterations in Fkbp10-/-mice. Hum. Mol. Genet. 2014;23:4822–31.

[10] Nigam SK, Jin YJ, Jin MJ, Bush KT, Bierer BE, Burakoff SJ. Localization of the FK506-binding protein, FKBP 13, to the lumen of the endoplasmic reticulum. Biochem. J. 1993;294:511–15.

[11] Bush KT, Hendrickson BA, Nigam SK. Induction of the FK506-binding protein, FKBP13, under conditions which misfold proteins in the endoplasmic reticulum. Biochem. J. 1994;303:705–08.

[12] Partaledis JA, Berlin V. The FKB2 gene of Saccharomyces cerevisiae, encoding the immunosuppressant-binding protein FKBP-13, is regulated in response to accumulation of unfolded proteins in the endoplasmic reticulum. Proc. Natl. Acad. Sci. U. S. A. 1993;90:5450–54.

[13] Boon K, Bailey NW, Yang J, Steel MP, Groshong S, Kervitsky D, et al. Molecular Phenotypes Distinguish Patients with Relatively Stable from Progressive Idiopathic Pulmonary Fibrosis (IPF). PLoS ONE. 2009. doi: 10.1371/journal.pone.0005134.

[14] Horimasu Y, Ishikawa N, Iwamoto H, Ohshimo S, Hamada H, Hattori N, et al. Clinical and molecular features of rapidly progressive chronic hypersensitivity pneumonitis. Sarcoidosis Vasc. Diffuse Lung Dis. 2017;34:48–57.

[15] Ishikawa Y, Mizuno K, Bächinger HP. Ziploc-ing the structure 2.0: Endoplasmic reticulum-resident peptidyl prolyl isomerases show different activities toward hydroxyproline. J. Biol. Chem. 2017;292:9273–82.

[16] Jeong M, Jang E, Choi SS, Ji C, Lee K, Youn J. The Function of FK506-Binding Protein 13 in Protein Quality Control Protects Plasma Cells from Endoplasmic Reticulum Stress-Associated Apoptosis. Front. Immunol. 2017. doi: 10.3389/fimmu.2017.00222.

[17] Ayaub EA, Kolb PS, Mohammed□Ali Z, Tat V, Murphy J, Bellaye P-S, et al. GRP78 and CHOP modulate macrophage apoptosis and the development of bleomycin-induced pulmonary fibrosis. J. Pathol. 2016;239:411–25.

[18] Vandesompele J, De Preter K, Pattyn F, Poppe B, Van Roy N, De Paepe A, et al. Accurate normalization of real-time quantitative RT-PCR data by geometric averaging of multiple internal control genes. Genome Biol. 2002;3:RESEARCH0034.

[19] Reyfman PA, Walter JM, Joshi N, Anekalla KR, McQuattie-Pimentel AC, Chiu S, et al. Single-Cell Transcriptomic Analysis of Human Lung Provides Insights into the Pathobiology of Pulmonary Fibrosis. Am. J. Respir. Crit. Care Med. 2018. doi: 10.1164/rccm.201712-2410OC.

[20] Butler A, Hoffman P, Smibert P, Papalexi E, Satija R. Integrating single-cell transcriptomic data across different conditions, technologies, and species. Nat. Biotechnol. 2018;36:411–20.

[21] Korfei M, Ruppert C, Mahavadi P, Henneke I, Markart P, Koch M, et al. Epithelial Endoplasmic Reticulum Stress and Apoptosis in Sporadic Idiopathic Pulmonary Fibrosis. Am. J. Respir. Crit. Care Med. 2008;178:838–46.

[22] Cabrera S, Selman M, Lonzano-Bolaños A, Konishi K, Richards TJ, Kaminski N, et al. Gene expression profiles reveal molecular mechanisms involved in the progression and resolution of bleomycin-induced lung fibrosis. Am. J. Physiol. - Lung Cell. Mol. Physiol. 2013;304:L593–601.

[23] Plataki M, Koutsopoulos AV, Darivianaki K, Delides G, Siafakas NM, Bouros D. Expression of Apoptotic and Antiapoptotic Markers in Epithelial Cells in Idiopathic Pulmonary Fibrosis. Chest. 2005;127:266–74.

[24] Thamsen M, Ghosh R, Auyeung VC, Brumwell A, Chapman HA, Backes BJ, et al. Small molecule inhibition of IRE1α kinase/RNase has anti-fibrotic effects in the lung. PLoS ONE. 2019. doi: 10.1371/journal.pone.0209824.

[25] Hsu H-S, Liu C-C, Lin J-H, Hsu T-W, Hsu J-W, Su K, et al. Involvement of ER stress, PI3K/AKT activation, and lung fibroblast proliferation in bleomycin-induced pulmonary fibrosis. Sci. Rep. 2017. doi: 10.1038/s41598-017-14612-5.

[26] Sriburi R, Bommiasamy H, Buldak GL, Robbins GR, Frank M, Jackowski S, et al. Coordinate Regulation of Phospholipid Biosynthesis and Secretory Pathway Gene Expression in XBP-1(S)-induced Endoplasmic Reticulum Biogenesis. J. Biol. Chem. 2007;282:7024–34.

[27] Zimmerman KA, Graham LV, Pallero MA, Murphy-Ullrich JE. Calreticulin Regulates Transforming Growth Factor-β-stimulated Extracellular Matrix Production. J. Biol. Chem. 2013;288:14584–98.

[28] Otsuka M, Shiratori M, Chiba H, Kuronuma K, Sato Y, Niitsu Y, et al. Treatment of pulmonary fibrosis with siRNA against a collagen-specific chaperone HSP47 in vitamin A-coupled liposomes. Exp. Lung Res. 2017;43:271–82.

[29] Tanaka K-I, Tanaka Y, Namba T, Azuma A, Mizushima T. Heat shock protein 70 protects against bleomycin-induced pulmonary fibrosis in mice. Biochem. Pharmacol. 2010;80:920–31.

[30] Moeller A, Ask K, Warburton D, Gauldie J, Kolb M. The bleomycin animal model: A useful tool to investigate treatment options for idiopathic pulmonary fibrosis? Int. J. Biochem. Cell Biol. 2008;40:362–82.

[31] Polosukhin VV, Degryse AL, Newcomb DC, Jones BR, Ware LB, Lee JW, et al. Intratracheal Bleomycin Causes Airway Remodeling and Airflow Obstruction in Mice. Exp. Lung Res. 2012;38:135–46.

[32] Uhal BD, Joshi I, Hughes WF, Ramos C, Pardo A, Selman M. Alveolar epithelial cell death adjacent to underlying myofibroblasts in advanced fibrotic human lung. Am. J. Physiol.-Lung Cell. Mol. Physiol. 1998;275:L1192–99.

[33] Kamp DW, Liu G, Cheresh P, Kim S-J, Mueller A, Lam AP, et al. Asbestos-Induced Alveolar Epithelial Cell Apoptosis. The Role of Endoplasmic Reticulum Stress Response. Am. J. Respir. Cell Mol. Biol. 2013;49:892–901.

[34] Nguyen H, Uhal BD. The unfolded protein response controls ER stress-induced apoptosis of lung epithelial cells through angiotensin generation. Am. J. Physiol.-Lung Cell. Mol. Physiol. 2016;311:L846–54.

[35] Nita I, Hostettler K, Tamo L, Medová M, Bombaci G, Zhong J, et al. Hepatocyte growth factor secreted by bone marrow stem cell reduce ER stress and improves repair in alveolar epithelial II cells. Sci. Rep. 2017. doi: 10.1038/srep41901.

[36] Ye R, Jung DY, Jun JY, Li J, Luo S, Ko HJ, et al. Grp78 heterozygosity promotes adaptive unfolded protein response and attenuates diet-induced obesity and insulin resistance. Diabetes. 2010;59:6–16.

[37] Ayaub EA, Dubey A, Imani J, Botelho F, Kolb MRJ, Richards CD, et al. Overexpression of OSM and IL-6 impacts the polarization of pro-fibrotic macrophages and the development of bleomycin-induced lung fibrosis. Sci. Rep. 2017;7:13281.

[38] Weiner SM, Sellin L, Vonend O, Schenker P, Buchner NJ, Flecken M, et al. Pneumonitis associated with sirolimus: clinical characteristics, risk factors and outcome—a single-centre experience and review of the literature. Nephrol. Dial. Transplant. 2007;22:3631–37.

[39] Morelon E, Stern M, Israël-Biet D, Correas JM, Danel C, Mamzer-Bruneel MF, et al. Characteristics of sirolimus-associated interstitial pneumonitis in renal transplant patients. Transplantation. 2001;72:787–90.

[40] Witt LJ, Demchuk C, Curran JJ, Strek ME. Benefit of adjunctive tacrolimus in connective tissue disease-interstitial lung disease. Pulm. Pharmacol. Ther. 2016;36:46–52.

[41] Kurita T, Yasuda S, Oba K, Odani T, Kono M, Otomo K, et al. The efficacy of tacrolimus in patients with interstitial lung diseases complicated with polymyositis or dermatomyositis. Rheumatol. Oxf. Engl. 2015;54:39–44.

[42] Barba T, Fort R, Cottin V, Provencher S, Durieu I, Jardel S, et al. Treatment of idiopathic inflammatory myositis associated interstitial lung disease: A systematic review and meta-analysis. Autoimmun. Rev. 2019;18:113–22.

[43] Koike R, Tanaka M, Komano Y, Sakai F, Sugiyama H, Nanki T, et al. Tacrolimus-induced pulmonary injury in rheumatoid arthritis patients. Pulm. Pharmacol. Ther. 2011;24:401–06.

[44] Sasaki T, Nakamura W, Inokuma S, Matsubara E. Characteristic features of tacrolimus-induced lung disease in rheumatoid arthritis patients. Clin. Rheumatol. 2016;35:541–45.

[45] Han Q, Lin L, Zhao B, Wang N, Liu X. Inhibition of mTOR ameliorates bleomycin-induced pulmonary fibrosis by regulating epithelial-mesenchymal transition. Biochem. Biophys. Res. Commun. 2018;500:839–45.

[46] Madala SK, Maxfield MD, Davidson CR, Schmidt SM, Garry D, Ikegami M, et al. Rapamycin Regulates Bleomycin-Induced Lung Damage in SP-C-Deficient Mice. Pulm. Med. 2011. doi: 10.1155/2011/653524.

[47] Lawrence J, Nho R. The Role of the Mammalian Target of Rapamycin (mTOR) in Pulmonary Fibrosis. Int. J. Mol. Sci. 2018. doi: 10.3390/ijms19030778.

