## Supplemental figures for "FK506-binding protein 13 expression is upregulated in interstitial lung disease and correlated with clinical severity: a potentially protective role"

**Supplementary Figure Legends**

**Supplemental Figure 1. Single-cell expression profile of FKBP13 expression in human IPF and control lung tissues.** The open access single-cell RNAseq dataset by Reyfman et al. was used to assess FKBP13 RNA expression in distinct cellular populations. Data from 8 healthy controls and 4 IPF patients were included in the analysis. (A) Cells were assigned to clusters based on predefined canonical expression markers. (B) Feature plot demonstrating FKBP13 expression in various cell populations. (C) Violin plot demonstrating FKBP13 mRNA expression in various cell populations from pooled IPF and control single-cell RNAseq data. *, $p < 0.05$ vs. healthy control.

**Supplemental Figure 2. Pressure-volume loops for bleomycin-treated mice.** (A,B) Pressure-driven pressure–volume (PV) loops was performed using flexiVent®. The average value of the animals in each group is shown. WT mice were unaffected with respect to lung function and histopathology, while FKBP13$^{-/-}$ mice displayed decreased static lung compliance. At the higher dose of bleomycin (0.06U), both WT and FKPB13$^{-/-}$ mice experienced a similar reduction in lung compliance at Day 21.

### Supplemental Figure 1

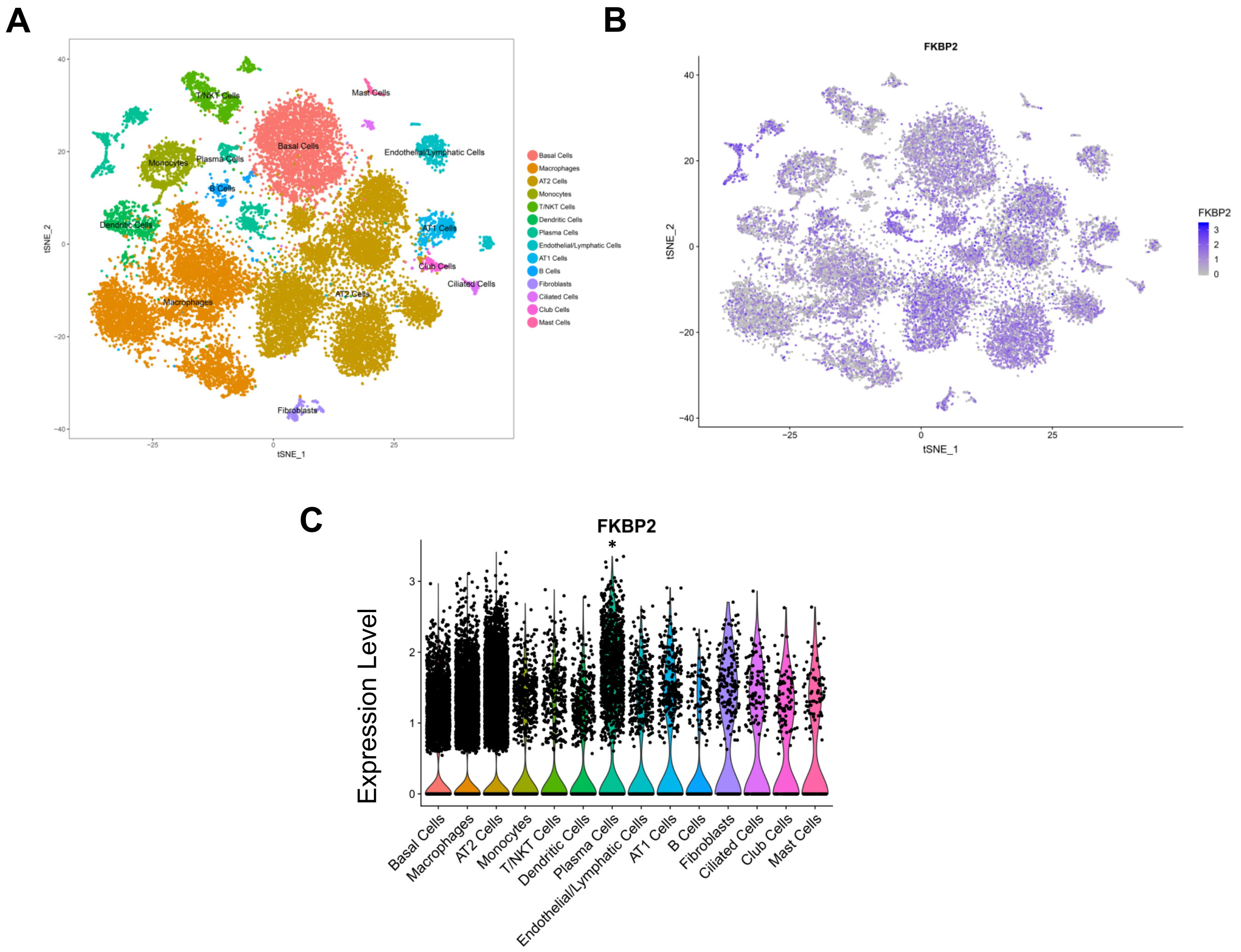

### Supplemental Figure 2

A

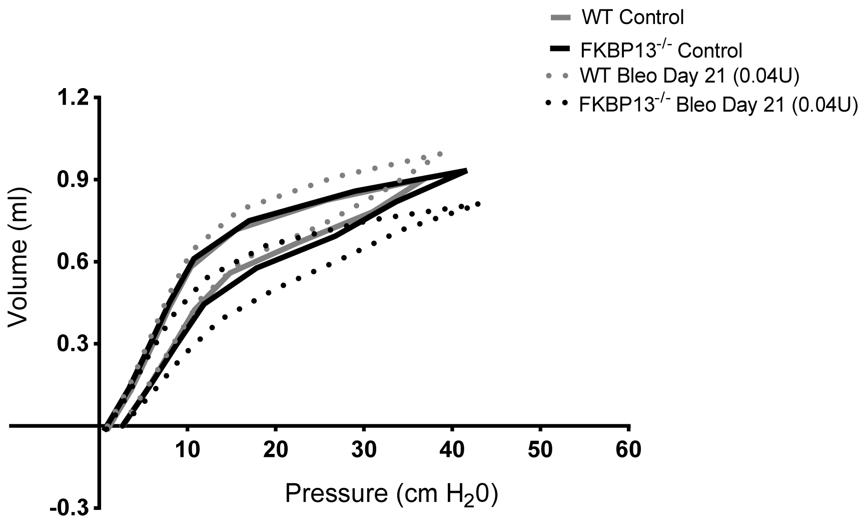


B

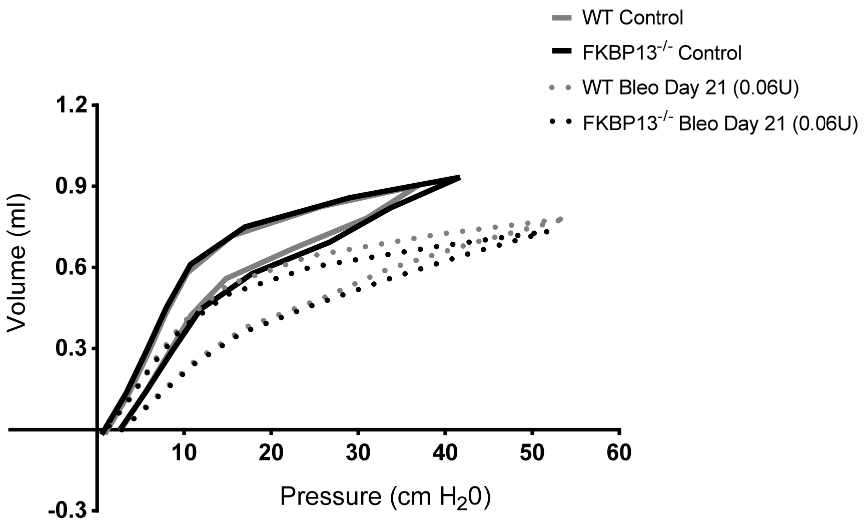
